# Quantitative Assessment of Colorectal Cancer Progression: a Comparative Study of Linear and Nonlinear Microscopy Techniques

**DOI:** 10.1101/398719

**Authors:** J. Adur, L. Erbes, M. Bianchi, S. Ruff, A. Zeitoune, M.F. Izaguirre, C.L. Cesar, H.F. Carvahlo, V.H. Casco

**Affiliations:** Instituto de Investigación y Desarrollo en Bioingeniería y Bioinformática (IBB), CONICET-UNER, Argentina.; Microscopy Laboratory Applied to Molecular and Cellular Studies, School of Bioengineering, National University of Entre Ríos, Argentina.; Department of Physics of Federal University of Ceara (UFC), Brazil.; INFABiC-National Institute of Science and Technology on Photonics Applied to Cell Biology, Campinas, Brazil.; Department of Structural and Functional Biology, Biology Institute, State University of Campinas, Brazil.

**Keywords:** colorectal cancer, linear microscopy, nonlinear microscopy, early diagnosis

## Abstract

**BACKGROUND AND AIMS:** Colorectal cancer (CRC) is a disease that can be prevented if is diagnosed and treated at pre-invasive stages. Thus, the monitoring of colonic cancer progression can improve the early diagnosis and detection of malignant lesions in the colon. This monitoring should be performed with appropriate image techniques and be accompanied by proper quantification to minimize subjectivity. We have monitored the mice CRC progression by image deconvolution, two-photon emission fluorescence (TPEF) and second harmonic generation (SHG) microscopies and present different quantization indices for diagnosis.

**METHODS:** The Azoxymethane (AOM) / dextran sodium sulfate (DSS) protocol was used. 35 eight-week old male BALB/cCmedc mice were used and distal colon segments were dissected at day zero and fourth, eighth, sixteen, and twenty weeks after injection. These segments were observed with linear and nonlinear optical microscopies and several parameters were used for quantification.

**RESULTS:** Crypt diameter higher than 0.08 mm and increased fluorescence signal intensity in linear images; as well as aspect relation above 0.7 and altered organization reflexed by high-energy values obtained from SHG images, away from those obtained in normal tissues.

**CONCLUSION:** The combination of linear and nonlinear signals improve the detection and classification of pathological changes in crypt morphology/distribution and collagen fiber structure/arrangement. In combination with standard screening approaches for CRC, the proposed methods improve the detection of the disease in its early stages, thereby increasing the chances of successful treatment.

## INTRODUCTION

Colorectal cancer (CRC) is a common malignancy, it is the third most common cause of cancer death in both men and women [1]. In general, CRC has a good prognosis, when it is diagnosed and treated at pre-invasive stages [2]. However, the detection of early lesions is still challenging.

To systematically characterize early tumoral transformations, an animal model of CRC progression is needed. Mice Azoxymethane (AOM)-induced CRC is one of the most commonly used pre-clinical animal model to mimic human sporadic CRC development [3]. Specifically, the AOM/dextran sodium sulfate (DSS) protocol has been proven a powerful tool for investigating the pathogenesis and chemoprevention of colitis-related colon carcinogenesis [4]. The development of colonic cancer is a multi-stage process [5] and the monitoring of colonic cancer progression can improve the early diagnosis and detection of malignant colon lesions. This examining should be performed with appropriate image techniques and be accompanied by proper quantification to minimize subjectivity.

Currently, anatomo-pathological examination is the diagnostic gold standard. The process involves many steps, prior to examination. It is time-consuming and expensive, and the excision itself has the inherent risks of bleeding and perforation. Thus, new imaging technologies might be tremendously helpful. Many examination techniques, such as computed tomography (CT), positron emission tomography (PET), magnetic resonance imaging (MRI), endorectal ultrasound (ERUS) lack the needed resolution or require exogenous contrasting agents [6]. Confocal endomicroscopy can reveal histological details during ongoing endoscopy. However, the usefulness of this imaging modality in patients is limited because of the need of fluorescent dyes [7]. Nonlinear and linear microscopy techniques provide unprecedented information that might help to overcome these limitations. Two-photon excitation fluorescence (TPEF), second harmonic generation (SHG), third harmonic generation (THG), and coherent anti-Stokes Raman scattering (CARS) microscopy exhibit several diagnostic advantages. These techniques are label-free, offer inherent three-dimensional resolution, use near-IR excitation resulting in superior optical penetration, low photodamage, and provide quantitative information [8]. In the last years, the nonlinear techniques have proven to be efficient for the diagnosis of CRC [9, 10]. Similarly, confocal or deconvolution microscopy, with optical sectioning capacity and exploring tissue autofluorescence (i.e. label free) were also used for quantitative and diagnostic purposes [11, 12].

Contrast mechanisms in conventional optical imaging are based on absorption, reflection, scattering and fluorescence, and the response recorded is linearly dependent on the intensity of the incident light (mercury lamp or continuous wave [CW] laser). In contrast, nonlinear optical effects used as the imaging contrast, occur when a biological tissue interacts with an intense laser beam (pulsed laser) and exhibits a nonlinear response to the applied field strength [13] (Fig. 1). Linear technique is cheaper and easily to operate while nonlinear needs specific and expensive instrumentation and qualified operator. Nonlinear optic presents better tissue penetration due to reduced scattering of larger wavelengths. However, both tools could be useful for the monitoring of CRC progression.

**Fig 1:**
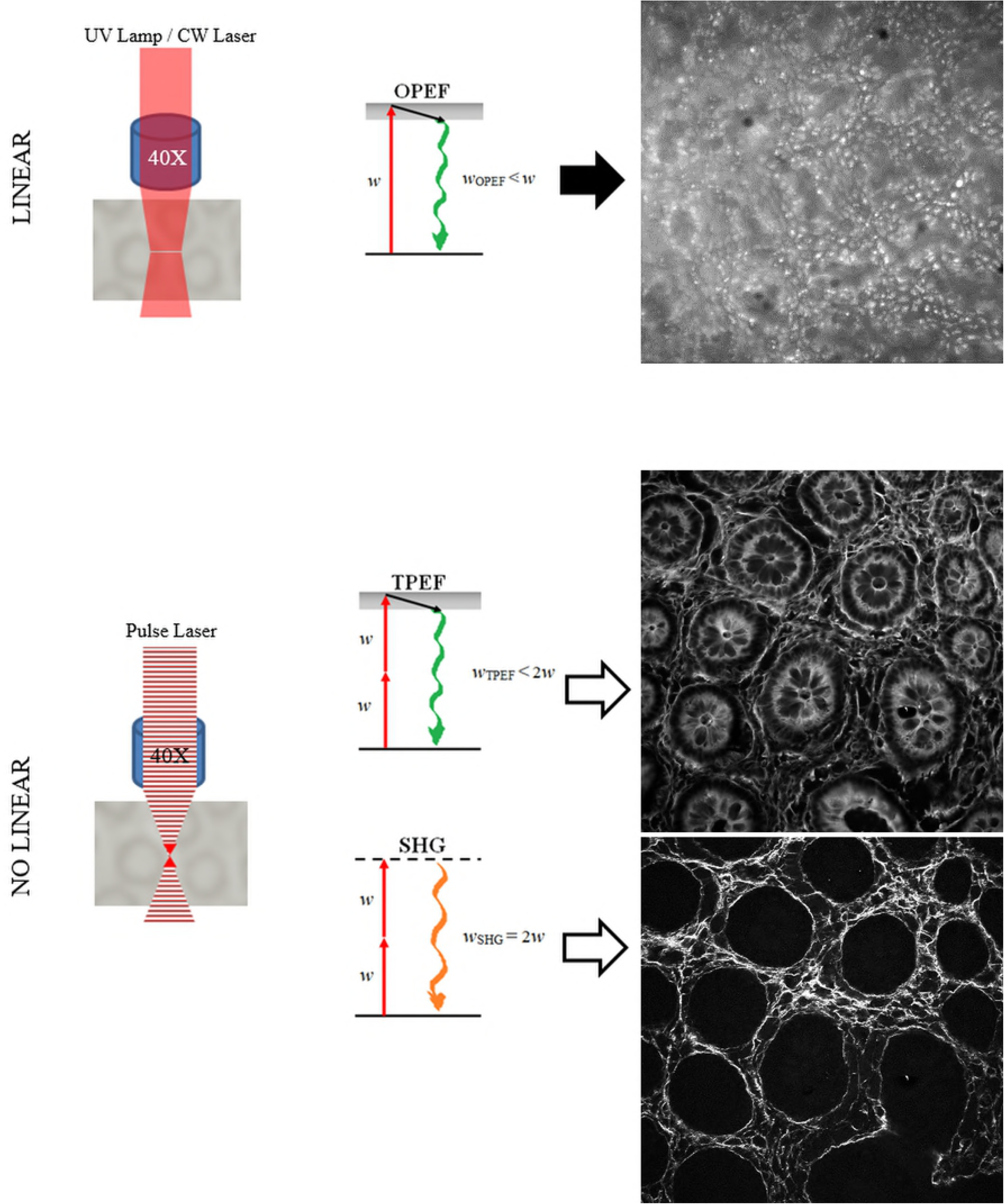
Schematic representation and Jablonski diagram of linear (one-photon emission fluorescence) and nonlinear (two-photon emission fluorescence and second harmonic generation) microscopy techniques (left column). Representative free-label (not stained) images of the mouse colon mucosa obtained respectively with these microscopy techniques are shown (right column).

In this work, we have examined CRC progression by image deconvolution, TPEF and SHG microscopies and present a detailed evaluation of the tumoral transforming colonic mucosa. We introduce the principle of operation and properties of each microscopy and present different quantization indices for diagnosis, discussing in depth the application of these novel techniques in a mouse model of CRC.

## MATERIALS AND METHODS

### -Murine model of AOM/DSS-induced colon cancer

The study was previously approved by the Ethics Committee for Animal Research of the National University of Entre Ríos. The use of laboratory animals followed the ethical code of International Organization of Medical Sciences for animal’s experimentation. The experimental population consisted of 35 eight-week old male BALB/cCmedc mice weighing 20g-30g. After 7 days of acclimatization, animals were randomized into experimental (AOM/DSS-treated) and control (saline-injected) groups. AOM/DSS-treated animals were intraperitoneally injected with azoxymethane (AOM) 0.01 mg/g body weight. One week later, they were given dextran sodium sulphate (DSS) in the drinking water for seven days, according to Tanaka and co-workers [14]. Distal colon segments were dissected at day zero and fourth, eighth, sixteen, and twenty weeks after injection (AOM-injected mice were not evaluated at the week 0) (Fig. 2A). These segments were destined to histological processing (for control purposes, Fig. 2B) and linear (Fig. 2C) and nonlinear optical (Fig. 2D) microscopy observation (for quantification).

**Fig 2:**
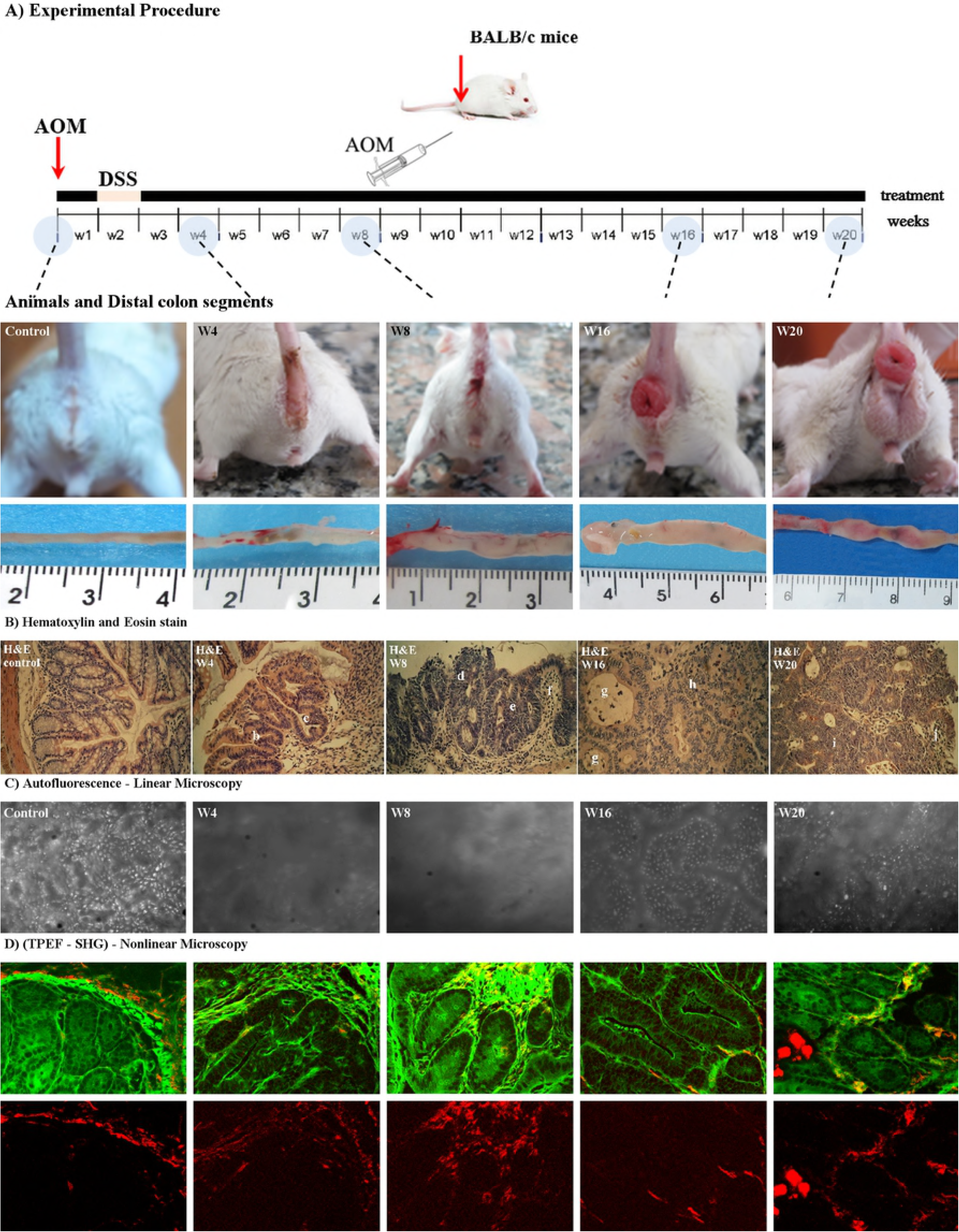
Experimental setup of mouse-CRC chemical induction. A) Mice were injected with AOM or saline solution (control) at day 0, one week later, DSS was supplied in drink water during seven days. Next animals were sacrificed at weeks 4th, 8th, 16th, and 20th (W4, W8, W16, W20). Top row pictures show the representative appearance of the anal region of control and AOM/DSS treated mice. Below is the row in which distal colon segments of control and CRC progression stages are shown. B) Representative H&E-stain sections used as gold standard. a: shortened crypts, b: bifurcated crypts, c: fused crypts, d: severe dysplasia, e: goblet cell depletion, f: lymphocytic infiltrate, g: dirty necrosis zone, h: absence of goblet cells, i: severe dysplasia, j: lymphocytic infiltrate. C) Representative images obtained with linear microscopy, and D) Representative non-linear microscopy: green (TPEF) and red (SHG) signals, respectively.

### -Linear Microscopy

A fluorescence deconvolution microscopy system was used as linear setup. This technique included: (a) optical-sectioning of the autofluorescent distal colon segments (producing a 3D image stacks), (b) Point Spread Function (PSF) determination, (c) deconvolution of the 3D image stacks, (d) 2D or 3D quantifications. The system used was based on an Olympus BX50 upright microscope; a complete description of the system has been previously published [15] (Fig. S1A, supplementary material). Deconvolution was performed using the DeconvolutionLab plugin from ImageJ [16]. Colon mucosa auto-fluorescence was observed at different wavelengths (Fig. 2C). Table 1 lists the major components identified by the autofluorescence.

**Table 1:**
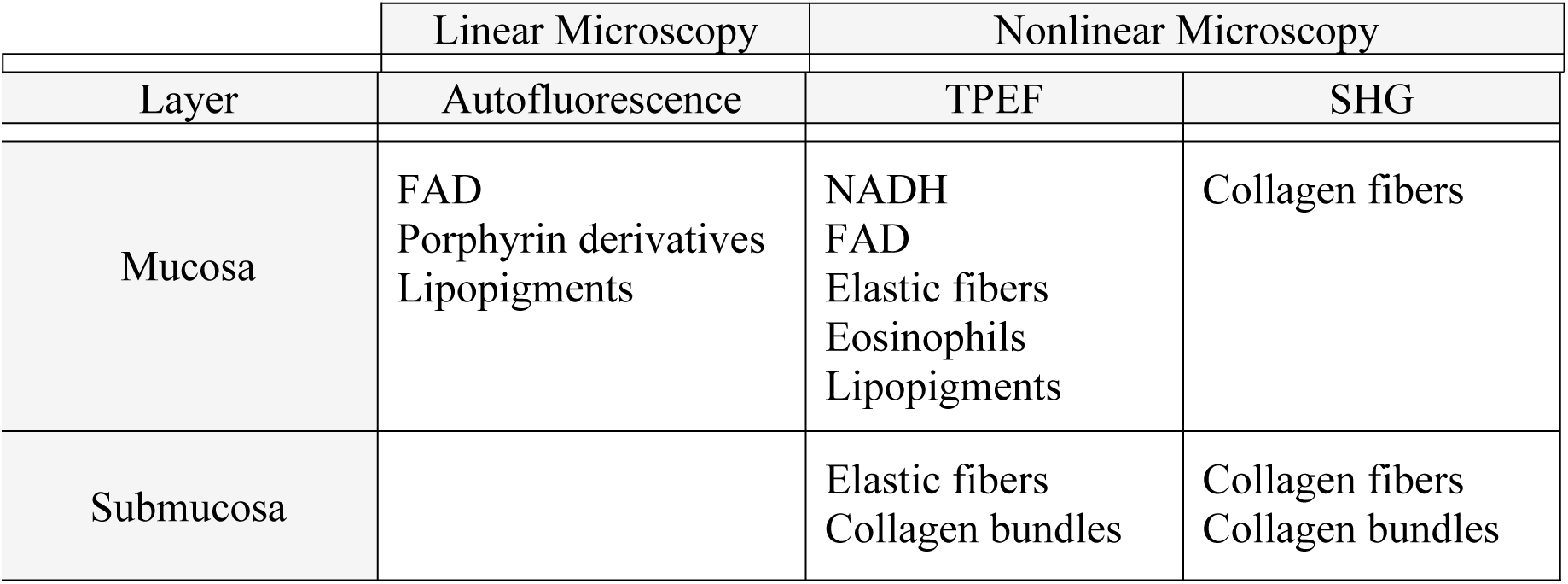
The origins of principal intrinsic signals in colon mouse

|  | Linear Microscopy | Nonlinear Microscopy |  |
| --- | --- | --- | --- |
| Layer | Autofluorescence | TPEF | SHG |
| Mucosa | FAD<br>Porphyrin derivatives<br>Lipopigments | NADH<br>FAD<br>Elastic fibers<br>Eosinophils<br>Lipopigments | Collagen fibers |
| Submucosa |  | Elastic fibers<br>Collagen bundles | Collagen fibers<br>Collagen bundles |

### -Nonlinear Microscopy

The system consisted of an inverted microscope Axio Observer Z.1 equipped with a Zeiss LSM 780-NLO confocal scan head (Carl Zeiss AG, Okerkochen, Germany). The excitation beam was provided by a mode-locked Ti:Sapphire laser (Spectra-Physics. Irvine, CA, USA) at the wavelength of 940 nm and an average power at the sample of 10 mW. The forward propagating SHG (λ=470 nm) and two-photon fluorescence signals (λ>490 nm) were acquired simultaneously by a non-descanned detector (NDD). A complete description of the system has been published elsewhere [17]. Figure 2D shows representative images acquired with this configuration; major components identified in this system are listed in Table 1. The microscope structure can be seen in Figure S1B (Supplementary material).

### -Acquisition Strategy

Murine colon segments were longitudinally splitted and fresh mucosa optically sectioned in apical-basal direction to acquire linear autofluorescence images (Fig. 3). Optical sectioning was performed to obtain tissue information through the crypt axis, up to 40 µm depth. After acquisition and before quantification, all images were deconvolved. Distal colon serial segments of each animal, in turn, were fixed, embedded on paraffin, sectioned to 5 µm thickness and stained with hematoxilin and eosin (HE) to classical histological analysis (methodological control). In addition, some sections were stained with the Schmorl and Fouchet stain [18] to detect lipofuscins and porphyrins. On the other hand, cross-sectioned murine colon segments were freshly observed to obtain nonlinear TPEF and SHG images (Fig. 3). Distal colon serial segments of each animal, in turn, were processed to histological analysis by HE-stain.

**Fig 3:**
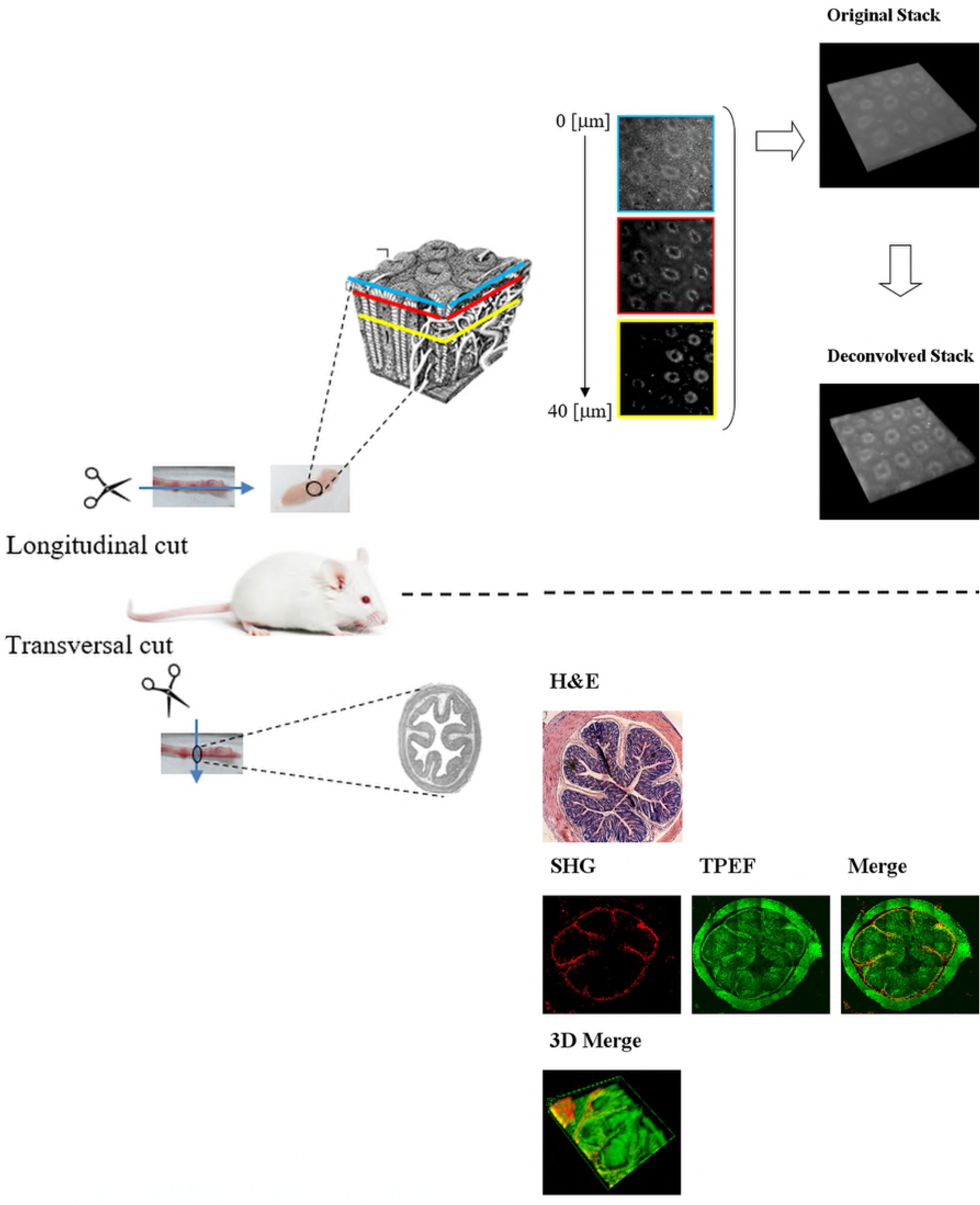
General methodological strategy: Top panel shows images acquisition from colon longitudinal sections. The open fresh mucosa is optically sectioned, and then deconvolved to obtain images at different depths; blue square: surface (depth 0), red square: superficial epithelium (5-10 µm depth) and yellow square: deep mucosa (30-40 µm depth). Bottom panel shows different representation possibilities of nonlinear TPEF (green) and SHG (red) images obtained from fresh tissue cross sections.

Three animals (saline-injected) were euthanized at time 0 and used as controls (time/week 0). The remaining 32 were euthanized at four, eight, sixteen and twenty weeks after the chemical injection saline or AOM. For each experimental point of AOM/DSS-treated animals, 8 mice were injected: 5 with AOM (colon cancer model) and 3 saline solution (control). For each colon segment, images from three different zones in the plane were captured in 2D studies and one sectioning of 40 µm (linear) or 100 µm (nonlinear) thickness for 3D analysis.

### -Quantification methodologies

The analysis of colon cancer progression was evaluated both at morphological and biochemical level by colon deconvolved autofluorescence images. For the morphological analysis was applied the Kudo’s pit pattern classification, to determine manual- and automatically, the shape and pit pattern of the colon crypt pits. Briefly, superficial colon crypts were categorized using Kudo’s classification, according to appearance, shape and perimeter [19]. For automatic analysis, 2D deconvolved images were segmented to identify crypt morphology. While Types I and II were considered benign changes, the pit pattern classes III-V were classified as neoplastic and malignant changes.

The biochemical analysis of the endogenous fluorophores changes was done for intensity measurements. Briefly, 2D images were opened in ImageJ. To avoid out-of-focus or noisy areas, a square template of 8×8 ROI (region of interest), each one of 64×64 pixels, was drawn and positioned on the image. Twenty ROIs were selected from the template to quantify the intensity. Using the multi-measure plugin, integrated density was measured for each ROI. Finally, considering a linear behavior of the CCD camera and the independence of the exposure time (t_exp_), the following formula was used to obtain the average gray level value (I): [I = (I_average =_ – I_background_)/t_exp_], where I_average_ is the integrated intensity measure, I_background_ correspond to intensity level register without sample, and t_exp_ is the capture time.

In parallel, a morphological analysis of the colon cancer progression also was done from nonlinear TPEF and SHG images. The methods of nonlinear image analysis were carried out with ImageJ software and calculations were made for 16 ROIs (256 × 256 pixels) chosen over selected images. For SHG intensity amount, signal was separated from the background with a threshold at level 50 from the 0 to 255 gray levels. From each ROI the area fraction (essentially a measure of SHG prevalence) was quantified. To achieve the aspect ratio (AR), Fast Fourier Transform (FFT) was used [20]. If the fibers have a parallel arrangement, the intensity plot of the FFT image, look as an ellipse and consequently will have a higher AR. In contrast, fiber with aleatory arrangement, exhibit an intensity plot of the FFT image, closer to the shape of a circle and with low AR. Texture features analysis, on the other hand, was based on the calculation of correlation (C) and energy (E) of the gray level co-occurrence matrix (GLCM) of the images [21]. The texture analyses were performed by GLCM-Texture plugin from Image-J, which was described by Walker and collaborators [22].

### -Statistics

For multi-group comparisons, one-way analysis of variance (ANOVA) and a post-hoc Tukey-Kramer test were used. In addition, t-testing was applied for two-group comparisons. The level of significance employed was significant (*) *p*<0.05 and very significant (**) *p*<0.01 and extremely significant (***) *p*<0.001. Data were analyzed with SPSS 10.0 software.

## RESULTS

### -CRC progression

Figure 2B shows the time staging of the mouse model of carcinogen-induced CRC progression. Figure S2 (Supplementary material) displays representative H&E-stained samples from control (a) and AOM-treated mice at weeks 4 (b), 8 (c), 16 (d) and 20 (e) after injection, as identified by an expert pathologist. In summary, normal animals presented crypts with small and uniformly distributed circular lumen on the mucosa surface (Fig. S2A). At week 4, the AOM-treated mice showed higher diameter crypts that lacked the uniform distribution of the controls. Also, tubular crypts and small aberrant crypt foci (ACF) were identified (Fig. S2B). At week 8, ACF were bigger and appeared near large Peyer´s Plates. Crypt distributions were not uniform, some with a large circular lumen and some with stellate pattern (Fig. S2C). At week 16, mouse exhibit big ACF and crypts with larger circular and stellate lumens. Tubular adenomas are also identified (Fig. S2D). Finally, at week 20 tubular adenomas and well-differentiated adenocarcinoma were observed. Large circular and small circular crypt lumens were observed (Fig. S2E).

### -Morphological and biochemical analysis during CCR progression

The changes on quantities and subtypes of tissular fluorophores make possible evaluate the development and progression of tumoral processes. Thus using excitation wavelengths between 460 and 550 nm, was possible to register mainly autofluorescence from flavins, porphyrins and lipopigments (Fig. S3, Supplementary material). This endogenous biomarkers were used to analyze crypts morphology and tissue function changes on the disease progression (Fig. 4, Fig. 6, and Fig. S4).

**Fig 4:**
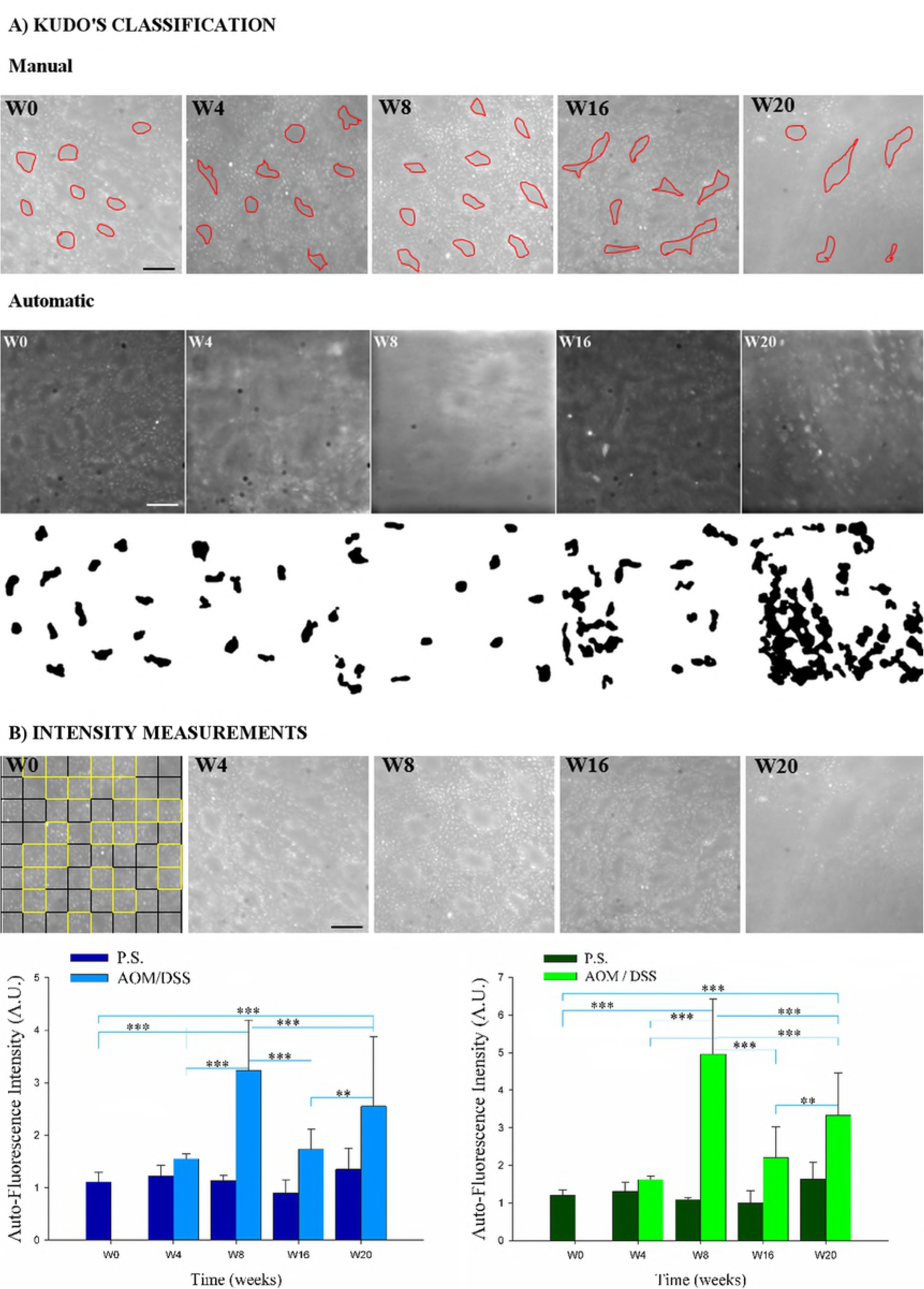
Autofluorescence images quantification. A) Morphological analysis using Kudo classification. First row shows representative images used to perform manual quantification, in which the red color lines (manually drawn), delimit the lumen of the crypts. Second row shows representative images used to perform automatic quantification. After segmentation method, individual crypts (black shapes) are measured. B) Intensity analysis of auto fluorescence images, exemplified in the first figure in which intensity variations (yellow squares) from a square template of 8×8 ROI, were evaluated. Blue and green bar graph shows intensity for two different filters. P.S: saline IP-injected animals and AOM/DSS: Azoxymethane and Dextran Sodium Sulfate treated animals.** indicates differences statistically significant (*p*< 0.05) and *** indicates differences statistically very significant (*p* < 0.01) following ANOVA. All scale bar = 30 [µm].

Figure 4 shows the results obtained by processing autofluorescence images. Figure 4A depicts images used for the Kudo’s pit pattern classification performed using manual (red lines) and automatic methodologies (black figures). Perimeters values were measured and the morphologies were identified with a help of a pathologist. A comparison table was made using perimeters values obtained from H&E-images, manual and automatic quantification data (Fig. S4, Supplementary material). Considering the appearance and perimeter of the crypt pits described by Hurlstone and Brown [23]; the Kudo’s pit pattern types were identified in the images used as samples. Through the manual methodology, benign changes such as normal round and star-shape pits (type I and II) were detected in time 0 of control samples (W0) and in all weeks of the treated samples (W4, W8, W16 and W20), while tubular pits (type IIIL) of neoplastic changes were also detected in the 16^th^ and 20^th^ weeks. The automatic methodology allowed to characterize more types of pits within the week analyzed and only 10.5% of pits were classified as “Possible turning to be Type II or IIIL” or “Possible turning to be IIIL” instead of being labelled with a particular Kudo’s pit pattern type. In control samples (with physiological solution), the 65% of the 276 quantified pits were identified as type I, whereas in treated samples, around 50% of the pits from the 4^th^ and 8^th^ pits were type I and the remaining were recognized as neoplastic changes (types III or IV). Data in Figure S4 (Supplementary material) evidence how averaged perimeter values for each period of time remained constant in control samples (0.06 mm) while they change in equivalent times of treated samples. Finally, the analyses of treated samples must be limited until the 8^th^week as the pits exhibit a progressive coalescence in the following weeks, which could lead to inaccurate identification and quantification of individual structures.

The analysis of autofluorescence intensity (Fig. 4B) show non-significant changes in the control animals (injected with saline) between weeks (*p* > 0.05). Intensity values range from 0.9 to 1.6 [arbitrary units, A.U.]. In AOM/DSS-treated animals, whereas the intensity increases slowly at week 4^th^(∼ 0.3 times), it shows a peak (∼ 3 times) at week 8^th^(differences statistically extremely significant *p* < 0.001). Striking, the endogenous fluorophores intensity increases ∼1.5 times at the week 16^th^and 2.5 times at the week 20^th^. The results found employing two different filters for excitation / emission respectively (460-490 nm / 515 nm; and 480-550 / 590 nm), show similar behavior.

### -Collagen analysis during CCR progression

The combination of TPEF (green) and SHG (red) images enables a superior evaluation of the spatial tissular organization during the disease progression, because the cellular and collagen extracellular matrix compartments configuration, respectively (Figures 5, 6, S5 and S6). Figure S5 (Supplementary material) displays the distribution of collagen fibers in representative mosaic images that allow to observe the complete colon cross-section. Changes in colon morphology (TPEF images) and collagen distribution (SHG images) were visualized during the disease progression. To confirm this preliminary data, collagen content both in mucosa and submucosa was quantified in each stage of cancer development (Fig. S6, Supplementary material). Collagen content significantly was significantly higher (*p*<0.01) in the submucosa as compared with mucosa at the different post-injection weeks. Interestingly, collagen content in the submucosa decayed significantly from time 0 (control) (64%) and week 4^th^ (16%), starting to recover from week 8^th^ (34%) and staying relatively constant (around 40%) in weeks 16^th^ and 20^th^. (Fig. S6B). To better understand the behavior of collagen fibers in the submucosa, additional parameters were quantified. Figure 5A shows representative submucosa regions where anisotropy (Fig. 5B), correlation (Fig. 5C) and energy (Fig. 5D) variables were measured. Figure 5B, represents the AR averages, at different disease stages. AR values significantly increase (*p* < 0.01) in diseased samples as compared to the normal colon along time. That means, fibers were arranged in parallel when compared to the AOM/DSS-treated mice samples, i.e. AR showed a more circular configuration (fiber without any specific arrangement). From texture analysis, fibrils correlation fell off sharply with distance at weeks 4^th^ and 8^th^(Fig. 5C), revealing the existence of isolated thin fibrils. On the other hand, correlation for the fibrils in normal and AOM/DSS-treated mice samples at weeks 16^th^ and 20^th^ remained elevated for larger distances, implying in more connected structures. Consistent with the qualitative observation, evaluation of three ROI (100×100 pixel side squares) in the SHG images showed that the correlation remained higher in the control and weeks 16^th^ and 20^th^; i.e. the Corr50 (the pixel distance where the correlation dropped below 50% of the initial value) significantly was greater in normal, W16, and W20 as compared with the early stages (W4 and W8) (Fig. 5C; *p*< 0.05, ANOVA). Since collagen fiber width and spacing affects the gray levels transition across the image, it is expected that GLCM energy will change for different collagen morphologies. Fibers in the normal submucosa have lower average energy because they are thinner, causing more variation in gray levels across image (Fig. 5D).

**Fig 5:**
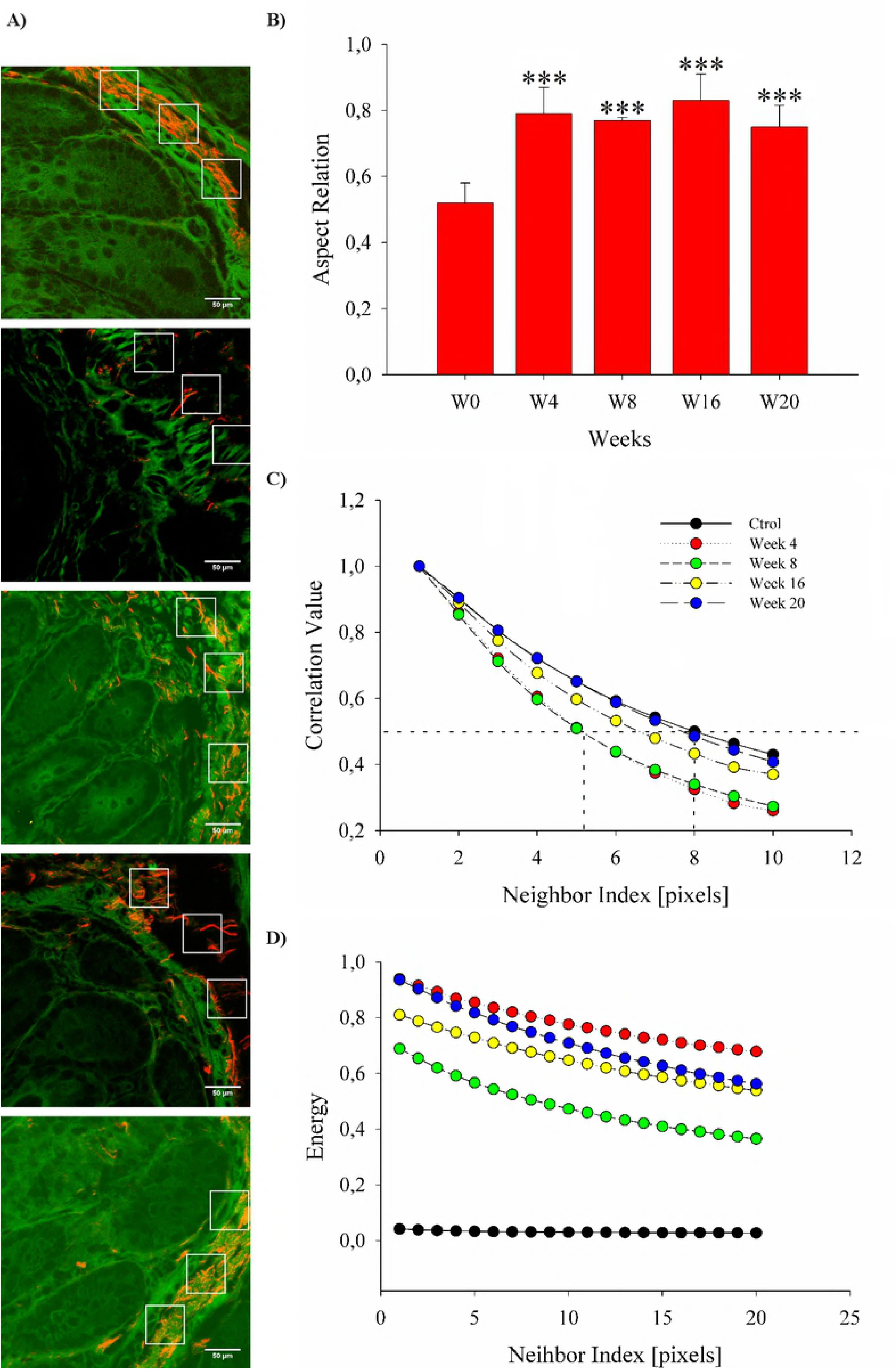
A) Submucosa TPEF+SHG images during CRC progression. B) Anisotropy calculation using aspect relation (AR). C) and D) texture parameters: correlation (C) and energy (D). Measurements were performed in 3 ROI of 100 × 100 pixel side squared (see white squared). *** indicates a statistically extremely significant (p < 0.01) difference. Scale bars = 50 [µm].

### -Analysis 3D of morphological and collagen transformations

Figure 6 shows representative 3D images allowing the qualitative analyzes of the auto-fluorescence and collagen arrangement. Original 3D images of the autofluorescence and deconvolved images (Figure 6A), revealed the morphology of the crypts. TPEF+SHG 3D images (Figure 6B) showed the variation in collagen fiber distribution around the crypts. The quantitative data obtained from the 2D images are qualitatively identified in the 3D aspects associated with CRC progression.

**Fig 6:**
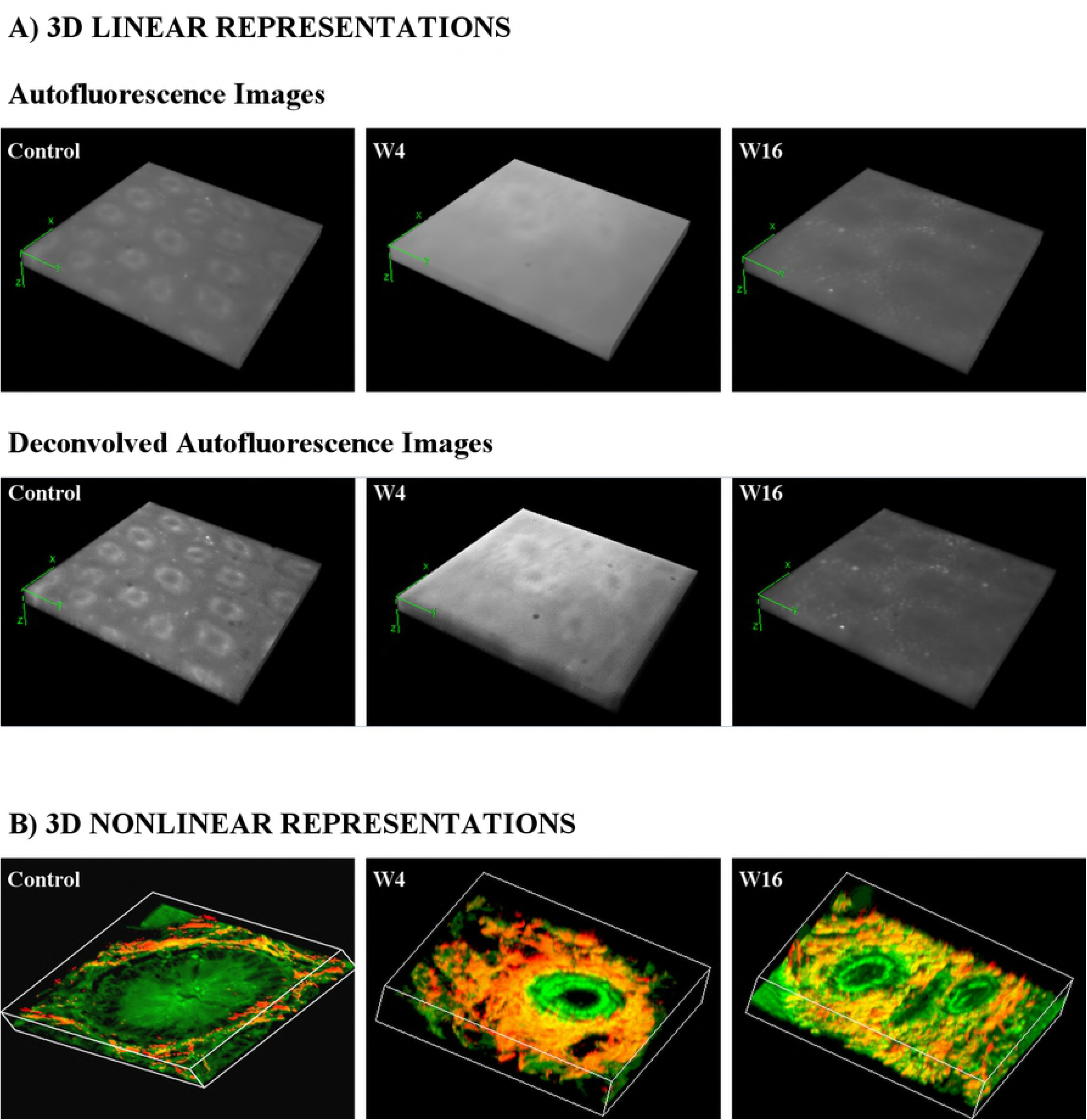
Representative 3D visualizations. A) Autofluorescence images before and after deconvolution process. Stack of 40 images, each separate 1 µm. B) Nonlinear collagen distribution (red). Stack of 100 images, each separate 1 µm.

## DISCUSSION AND CONCLUSION

Colon cancer is a disease with important repercussions on public health, its diagnosis and the early localization of cell/tissue transformations play an important role in the prevention and curative treatment of colonic cancer. However, physical biopsies do not fully solve this problem because are randomly sampled, and are highly dependent on the skill and experience of physician. One and two photon-induced autofluorescence depends on endogenous fluorophores of biological tissues, which undergo malignant transformation-related changes. Therefore, new techniques for the detection of pathological colonic tissue are needed. This report demonstrates that linear and nonlinear techniques could be used as effective methods for detecting and differentiating the abnormal tissues from the surrounding normal tissues. More importantly, the methods proposed here, not use exogenous markers to detect early changes, making them very promising for early diagnosis.

Linear signal was obtained using not-harmful 460 or 550 nm excitation wavelength range and allowed to precisely identify the lumen crypts of the colon mucosa. Endogenous markers showing fluorescence at these wavelengths were mostly flavins [24, 25], lipopigments, which exhibit bright dotted fluorescence [26, 27], and porphyrins [26-28]. The fluorescence images were used to characterize the crypts morphology and to determine variations in the autofluorescence intensity along the progression of colon cancer. Even though, several groups have reported the utility of autofluorescence to analyze colon tissues [11, 27, 29, 30], according our knowledge, this is the first time that this kind of signal is used to perform Kudo classification. Important, the Kudo´s classification differentiates neoplastic and non-neoplastic colon polyps, according to crypts morphology and its lumen size, employing a Narrow Band Image system (NBI) or chromoendoscopy with magnification. Based on Prieto et al., 2016 [30], the perimeter of the crypt lumens was measured and used for crypt pits classification by manual and automated analysis of autofluorescence microscopy images. Results show average perimeters of the distal colon crypt lumens coincident with mucosa benign changes (type I and II of Kudo’s pit pattern) at the week 4^th^ post-AOM/DSS-treatment [31]. Kudo´s work and other such as Tan et al., 2013 [32] were performed by images manual processing obtained from colon mucosa crypt pits with benign morpho-functional changes. Several analysis based on Kudo’s pit pattern classification in turn, have employed visual classifications by experienced endoscopists while just a few added an image processing software [30, 33] that work over chromoendoscopy [34, 35] and narrow band imaging endoscopy (NBI) images [36, 37]. The free-label method apply in the present work shows acceptable results in about 60 seconds, compared with Takemura´s procedure that take several minutes [33]. In contrast, the conventional procedure takes around 70 hours from tissue dissection to image reconstruction and analysis [38]. In addition to recognize the geometry of the crypts, endogenous fluorescent molecules allow to identify physiological and biochemical variations during tumoral disease evolution [39-41]. Thus, the fluorescence spectral features which differentiate normal mucosa and adenoma have been identified and empirically correlated with histological diagnosis. However, few works show changes in early stages as observed in this work. Thus, the animals induced to develop cancer presented an intensity peak (statistically extremely significant) at week 8^th^ post-AOM/DSS treatment. In general week 8^th^, 16^th^ and 20^th^ post-AOM/DSS treatment showed higher intensity values compared to the control and CCR early stage in week 4^th^. These results are coincident with previous reports [11, 27, 29] where an increase in autofluorescence signal is detected in dysplastic respect to normal tissues. Cellular autofluorescence is known to depend on the metabolic state [11], which changes not only through several physiological state but also pathological. According excitation wavelengths used in this work, the increased autofluorescence from week 4 of CCR induction could be due to hemoglobin breakdown products such as porphyrin derivatives, which are known to accumulate in tumor cells [28]. Porphyrins have excitation maxima near 400nm, and minor pick to 450-700 nm. Porphyrins also are localized on mitochondrial electron transport strand, highly developed during tumoral state to meet the metabolic demand [42]. Mitochondrias play critical roles in meeting cellular energy demand, in cell death, and in reactive oxygen species (ROS) and stress signaling [11].

Increasing autofluorescence, also can be attributed to lipofuscin and lipofuscin-like compounds dots, which increase under a variety of pathophysiological conditions [43, 44]. Between these, oxidative stress and ageing [45, 46], characterized by a progressive unbalance between protein damage and clearance, leading at its turn to an increased protein homeostasis disturbance, with accumulation of oxidized proteins’ aggregates and, subsequently, of highly-cross linked materials such as lipofuscin and lipofuscin-like lipopigments, affecting cell viability. These cytoplasmic aggregates of undigested cell materials resulting from phagocytosis and autophagy processes and accumulating as endocytoplasmic vesicles under both physiological and pathological conditions, such neoplastic state [47-49].

It is known that collagen networks of extracellular matrix is altered from several pathological states, like tumor development [50, 51]. Exploring harmless analysis techniques, recent studies that utilize SHG microscopy have discriminated alterations in collagen fiber scaffoldings associated with non-neoplastic [52, 53] and neoplastic diseases [9, 13, 54]. Such spatial alterations may provide cues to cell migration and invasion during cancer progression [55, 56]. Therefore, in order to evaluate the fibrillar networks 3D organization biological tissues, researchers have developed different metrics to quantify data from SHG images [56], such as fibers orientation, spacing, and thickness [57, 58]. Thus, using anisotropy, energy and aspect relation parameters of collagen fibers organization, in this work, the CRC progression was characterized. Fundamentally, the amount and reorientation of the collagen fibers decrease significantly and notably at week 4^th^, an early stage of colon cell-transformation. These results are coincident with our previous [9] and other works [59-61] that analyzed CRC. However, this is the first time that early changes in collagen network is demonstrated by several and harmless methods, and in a complete colon cancer progression. Both linear and nonlinear microscopies, using different strategies, showed that 3D-representations are extremely useful. With linear techniques, crypts transformations were visualized and modifications in autofluorescence intensity observed. In particular, 3D representations using nonlinear techniques allowed to clearly visualizing the modification and change of direction of the collagen fibers around the crypts. We believe that new algorithms are needed to extract quantitative parameters for these stacks.

In conclusion, linear and nonlinear signals improve the detection and classification of pathological changes in crypt morphology/distribution and collagen fiber structure/arrangement, and provide a more comprehensive understanding of the connection between CRC and tissue heterogeneity. In brief, crypt diameter higher than 0.08 mm and increased fluorescence signal intensity in linear images; and aspect relation above 0.7 and high-energy values obtained from SHG images indicate altered organization, disparate from that characteristic of the normal tissue. The independence of contrast agents arguably improve safety, cost, and time efficiency for diagnosis. In our opinion, technologies that identify reliable changes at the cellular level, characterizing the premalignant state, enable targeted biopsies and have the potential to dramatically reduce the number and cost of random biopsies. In combination with standard screening approaches for CRC, the proposed methods improve the detection of the disease in its early stages, thereby increasing the chances of successful treatment.

## ACKNOWLEDGEMENTS

J.A. is thankful to IBB-CONICET-UNER for funding (PIO Res: 4337/15; Contract grant number: N8 14620140100004 CO). H.F.C. is thankful to FAPESP for funding (2009/16150-6). INFABiC is co-funded by FAPESP (2014/50938-8) and CNPq (465699/2014-6).

## SUPLEMTARY FIGURE LEGENDS

**Figure S1:** Pictures of the microscopy systems used. A) The linear analysis was done using an epi-fluorescence microscopy Olympus BX50, adapted to perform optical sectioning and deconvolution methods. B) Nonlinear technique was based in the laser confocal Zeiss LSM780 and a pulsed Ti:Sapphire laser (Spectra-Physics. Irvine, USA) and OPO (APE, Levante, Berlin, Germany) system to perform CARS.

**Figure S2:** Histopathological CRC progression in mouse AOM/DSS model. S0) (a.1 and a.2): Longitudinal section of normal colon, the Lieberkühn crypts are observed as individual unbranched structures, with normal goblet cell content. S4) (b.1) Image of a pair of crypts with mild dysplasia are observed (arrows). (b.2 and b.3): Injuries known as “aberrant crypt foci” (arrows), in which, elongated, branched and fused crypts are observed. The aberrant crypt cells show hyperchromatic enlarged nuclei, with polarity loss. Also, can be observed a decrease in the goblet cell number. S8) (c.1-c.4): Microadenoma with adenocarcinoma in situ and abundant lymphocytic infiltrates. S16) (d.1): Severe dysplasia (arrow). (d.2-d.4): Intramucosal adenocarcinoma (arrows). S20) (e.1-e.4): Intramucosal adenocarcinoma, with abundant areas of necrotic debris (arrows).

**Figure S3:** H&E stain (Left column) and specific stain to detect porphyrin (Midle column) and lipofuscin (Right column) from representative images of CRC model.

**Figure S4:** Automatic quantification of autofluorescence images. Representative images at control, week 4^th^, 8^th^, 16^th^ and 20^th^ in A) untreated animals (saline injected) and B) AOM injected and DSS treated animals. Below each image the average perimeters of crypts are shown. These quantification and those made in H&E images are compared and presented in the table, where according to the morphology and perimeter crypts value are classified according Kudo’s criterion.

**Figure S5:** Representative mosaic images during the progression of CRC of a complete colon section showing morphological colon changes (TPEF-green) and collagen arrangement (SHG-red).

**Figure S6:** Collagen arrangement through the CRC progression in: A) submucosa colon regions. B) Quantification of collagen content in mucosa (images not show) and submucosa regions. For calculations 16 ROI (256 × 256 pixels) in each image were selected. Each bar represents the mean ± SD of three independent images of each point. Scale bar = 10 [µm].

